# Targetable cellular signaling events drive arterial rupture in knock-in mouse models of vascular Ehlers-Danlos Syndrome

**DOI:** 10.1101/627653

**Authors:** Caitlin J. Bowen, Juan Francisco Calderón Giadrosic, Graham Rykiel, Zachary Burger, Elaine C. Davis, Mark R. Helmers, Elena Gallo MacFarlane, Harry C. Dietz

**Affiliations:** Institute of Genetic Medicine, Johns Hopkins University School of Medicine, Baltimore, MD, USA 21287; Howard Hughes Medical Institute, Chevy Chase, MD, USA 20815; Anatomy and Cell Biology, McGill University, Montreal, Quebec, Canada H3A 0C7

## Introduction

Vascular Ehlers-Danlos Syndrome (vEDS) is an autosomal-dominant connective tissue disorder caused by heterozygous mutations in the COL3A1 gene^1^. Currently, loss of structural integrity of the extracellular matrix is believed to drive the signs and symptoms of this condition, including spontaneous arterial dissection and/or rupture, the major cause of mortality^2–4^.

Using novel mouse models of vEDS that carry heterozygous *Col3a1* glycine substitutions, we show that signaling abnormalities in the PLC/IP_3_/PKC/ERK pathway (Phospholipase C/Inositol 1,4,5-triphosphate/Protein Kinase C/Extracellular signal-regulated kinase) are major mediators of disease pathology and dissection. Treatment with pharmacologic agents that inhibit ERK1/2 and PKCβ activation prevent death due to spontaneous aortic rupture. Additionally, we find that pregnancy-or puberty-associated accentuation of vascular risk, also seen in vEDS patients, is rescued by attenuation of oxytocin or androgen signaling, respectively. Taken together, our results provide the first evidence that targetable signaling abnormalities contribute to the pathogenesis of vEDS, highlighting unanticipated therapeutic opportunities.

## Main

vEDS is associated with spontaneous rupture of any medium-to-large artery that often occurs without prior vessel enlargement^1–4^. Patients are also at risk for sudden rupture of hollow organs, including the bowel or gravid uterus^1,4^. The presenting signs in the majority of adults with vEDS are vascular dissection or organ rupture, with 25% of patients experiencing a major complication by 20 years of age^1,4^. Little is known about the pathogenesis of this disease and promising treatment strategies remain elusive^5–9^.

In order to investigate the mechanisms of aortic dissection, we used CRISPR/Cas9 to create two mouse models of vEDS, *Col3a1*^G209S/+^ and *Col3a1*^G938D/+^, which carry heterozygous knock-in glycine substitutions in critical residues involved in collagen triple helix assembly (Fig. 1A, B and Supplementary Table 1).

**Figure 1.**
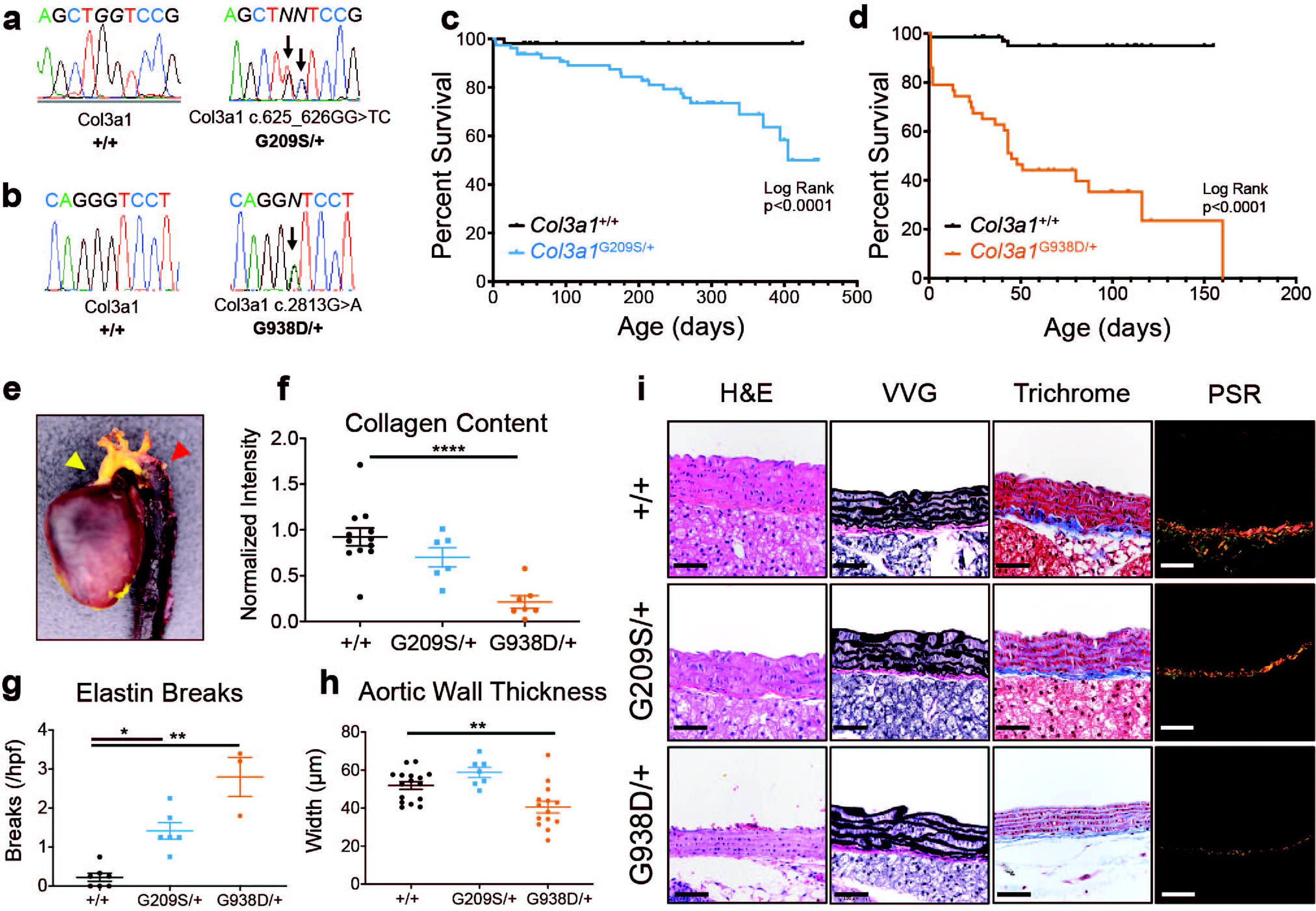
*Col3a1*^G209S/+^ and *Col3a1*^G938D/+^ mice recapitulate vEDS phenotypes. **a**, Sanger sequencing of genomic DNA confirmed the intended Col3a1 c.625_626GG>TC corresponding to G209S. **b**, Sanger sequencing of genomic DNA confirmed the intended Col3a1 c.2813G>A corresponding to G938D. **c**, Kaplan-Meier Survival curve for comparing *Col3a1*^+/+^ (n=53) to *Col3a1*^G209S/+^ mice (n=79), which die from vascular rupture or dissection. Significant differences were calculated using Log-Rank (Mantel-Cox) analysis. **d**, Kaplan-Meier Survival curve for comparing *Col3a1*^+/+^ (n=70) to *Col3a1*^G938D/+^ mice (n=43), which die from vascular rupture or dissection. Significant differences were calculated using Log-Rank (Mantel-Cox) analysis. **e**, A representative gross post-mortem examination of a latex-injected aorta (yellow arrowhead) demonstrates descending aortic dissection (red arrowhead). **f**, Quantification of collagen content in aortic cross sections, as measured by normalized PSR intensity. Error bars show mean ± s.e. Asterisks signify significant differences using one-way ANOVA with Dunnett’s multiple comparisons post-hoc test (****p<0.0001, DF = 2, F=13.97) **g**, Quantification of elastin breaks in aortic cross sections. Error bars show mean ± s.e. Asterisks signify significant differences using Kruskal-Wallis with Dunn’s multiple comparisons post-hoc test (*p<0.05, **p<0.01) **h**, Quantification of aortic wall thickness in aortic cross sections. Error bars show mean ± s.e. Asterisks signify significant differences using one-way ANOVA with Dunnett’s multiple comparisons post-hoc test (**p<0.01, DF = 2, F=10.16). **i**, Histological staining (H&E = Hematoxylin & Eosin, VVG = Verhoeff Van Gieson, Masson’s Trichrome, and PSR = Picrosirius Red) of wild type and vEDS aortic cross sections. Scale bar = 50 μm

Both the *Col3a1*^G209S/+^ and *Col3a1*^G938D/+^ mouse models recapitulate vEDS vascular phenotypes. Mice die suddenly due to aortic rupture, aortic dissection or organ rupture, presenting with hemothorax or hemoperitoneum at necropsy. *Col3a1*^G938D/+^ mice present with a more severe phenotype, with a median survival of 45 days compared to 400 days for the *Col3a1*^G209S/+^ mice, p<0.0001, Fig. 1C-E). Neither mouse model shows a tendency for formation of aortic root or ascending aortic aneurysm, consistent with human phenotypes (Supplementary Fig. 1). Although the aortic wall architecture is relatively preserved in both models, minor alterations include occasional elastic fiber breaks, decreased aortic wall thickness, and decreased collagen content (Fig. 1F-I). Analysis by transmission electron microscopy shows disruption of elastic lamellar units including thickened elastic fibers with a moth-eaten appearance, disarray of vascular smooth muscle cells (VSMCs) between fibers, and a paucity of collagen fibrils that normally occupy the intervening space between VSMCs and adjacent elastic fibers (Supplementary Fig. 2). Collagen fibrils within the aortic media showed wide variation in diameter with a generally smaller size when compared to control mice (Supplementary Fig. 2), consistent with previous reports^2,8,10^. Fibroblasts within the aortic adventitia of vEDS mice showed gross distension of the endoplasmic reticulum presumably due to impaired trafficking of abnormally folded type III collagen (Supplementary Fig. 2).

In order to evaluate the effect of drugs previously tested in other models of aortic disease such as Marfan (MFS) and Loeys-Dietz Syndrome (LDS)^11–16^, we assessed the effect of losartan, propranolol, atenolol and amlodipine on survival of vEDS mouse models. Despite all these drugs causing the predicted reduction in blood pressure (Supplementary Fig. 3), no drugs resulted in increased survival, with losartan, propranolol, and atenolol having no impact, and amlodipine actually increasing the risk of aortic dissection (Supplementary Fig. 3).

We also tested the effect of celiprolol, a β1 antagonist/β2 agonist that previous work has proposed, on the basis of a small trial, to delay adverse events in patients with vEDS^17,18^. Surprisingly, celiprolol, while having the predicted effect on pulse rate, accelerated death from aortic dissection in both the severe *Col3a1*^G938D/+^ and mild *Col3a1*^G209S/+^ vEDS mouse models (Supplementary Fig. 3). The fact that propranolol, a nonspecific β1/β2 antagonist, and atenolol, a specific β1 antagonist, had no impact on survival, suggest that this deleterious effect does not extend to all β-blockers and may be related to the drug’s β2 agonism and/or α2 antagonism^19^.

In order to elucidate which signaling abnormalities may mediate disease pathology in vEDS, we performed comparative transcriptional profiling by high-throughput RNA sequencing (RNA-seq) on the proximal descending thoracic aortas of *Col3a1*^G938D/+^ and *Col3a1*^+/+^ mice. The proximal descending thoracic aorta is where dissection is most commonly observed in these models.

We identified 170 consistently differentially expressed transcripts (probability of differential expression >0.95) between *Col3a1*^G938D/+^ aortas compared to *Col3a1*^+/+^ aortas (Fig. 2A). Network analysis indicated elevated mitogen-activated protein kinase (MAPK) activity, including p38, c-Jun N-terminal kinases (JNK), Akt, and extracellular signal–regulated kinases (ERK1/2) (Supplementary Fig. 4). Furthermore, upstream and gene set enrichment analyses suggested that transcriptional differences in vEDS aortas may be driven by excessive activity of ERK and Ca^++^/G_q_ protein coupled receptors (GPCRs), which signal through the PLC/IP_3_/PKC/ERK (phospholipase C/inositol 1,4,5-triphosphate/protein kinase C/extracellular signal-regulated kinase) axis (Fig. 2B).

**Figure 2.**
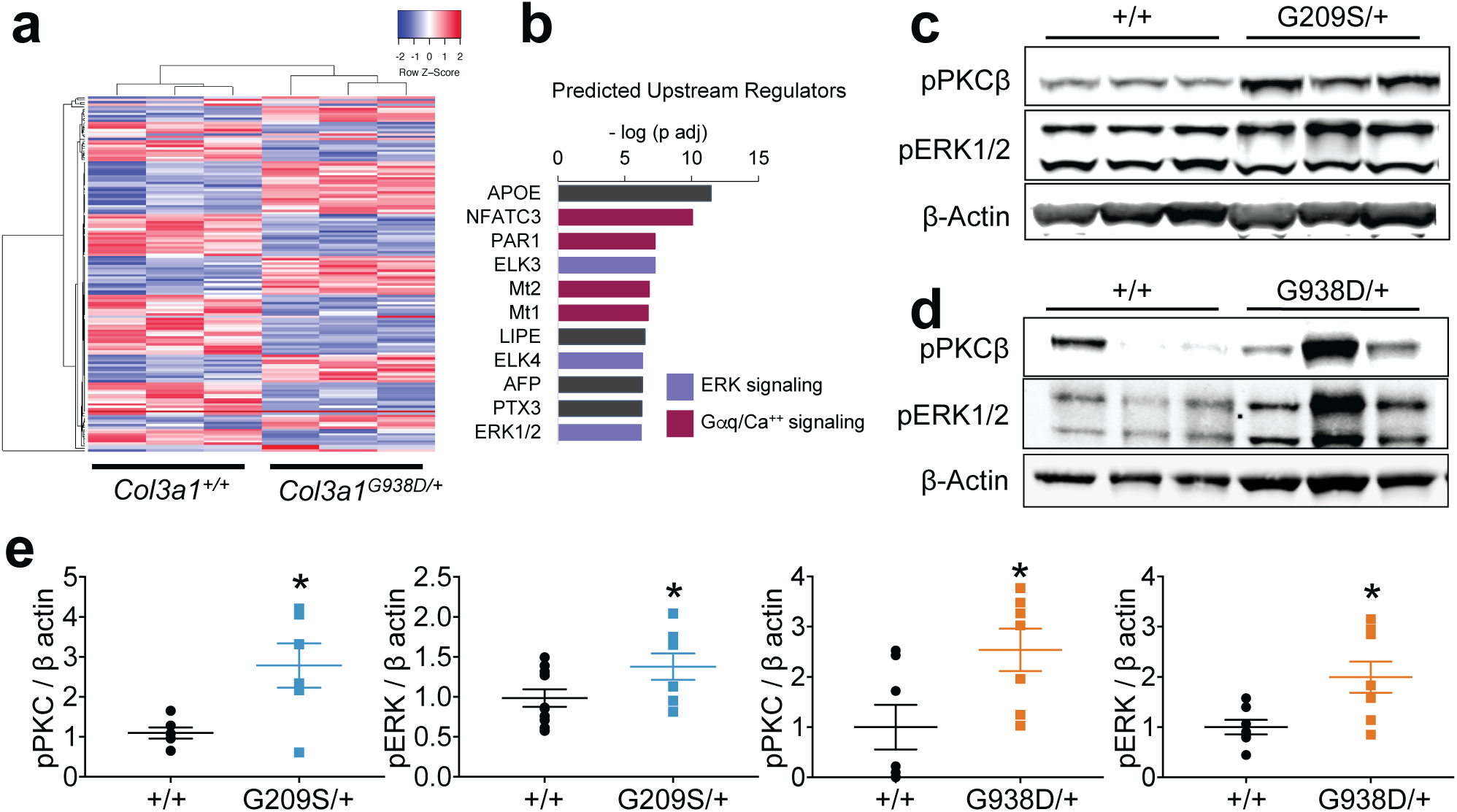
vEDS aortas display a molecular signature for excessive PKC/ERK signaling. **a**, Unsupervised hierarchical clustering using the most differentially expressed genes from bulk RNAseq was performed, and vEDS samples clustered separately from controls. **b**, Upstream analysis based on differentially expressed genes. Significant enrichment was determined using Fisher’s exact test. **c**, representative western blot analysis of pPKCβ and pERK comparing *Col3a1*^+/+^ to *Col3a1*^G209S/+^ proximal descending aortas. **d**, representative western blot analysis of pPKCβ and pERK comparing *Col3a1*^+/+^ to *Col3a1*^G938D/+^ proximal descending aortas. **e**, quantification of pPKCβ and pERK levels normalized to β-actin loading control comparing *Col3a1*^+/+^ (n=6) to *Col3a1*^G209S/+^ (n=6) and *Col3a1*^G938D/+^ (n=8) aortas. Error bars show mean ± s.e. Asterisks signify significant differences using two-tailed Student’s t-test (pERK/G209S T=2.053, DF=16; pPKC/G209S T=2.950, DF=10; pERK/G938D T=2.770, DF=13; *p<0.05) or Mann-Whitney test (G938D/pPKC; *p<0.05) depending on Shapiro-Wilk normality tests.

As predicted by transcriptome profiling, immunoblots of aortic lysates derived from control, *Col3a1*^G938D/+^ and *Col3a1*^G209S/+^ mice showed significantly increased ERK1/2 and PKCβ phosphorylation in the aortas of both vEDS mouse models (Fig. 2C-E). In view of the beneficial effects of PLC/IP_3_/PKC/ERK axis inhibition in mouse models of MFS^11^, we then tested if attenuation of this signaling pathway would decrease the risk of aortic rupture in vEDS mice. Treatment of *Col3a1*^G938D/+^ mice with ruboxistaurin, an orally administered pharmacologic agent that specifically inhibits PKCβ,^20^ resulted in 94% survival after 45 days of treatment, compared to only 55% survival with no treatment (Fig. 3A). As expected, inhibition of PKCβ with ruboxistaurin prevented autophosphorylation of PKCβ as well as phosphorylation of ERK1/2 in the aortic wall (Fig. 3B), as assessed by immunoblots of aortic lysates. These data suggest that PKC-dependent ERK activation is a critical driver of aortic disease in vEDS.

**Figure 3.**
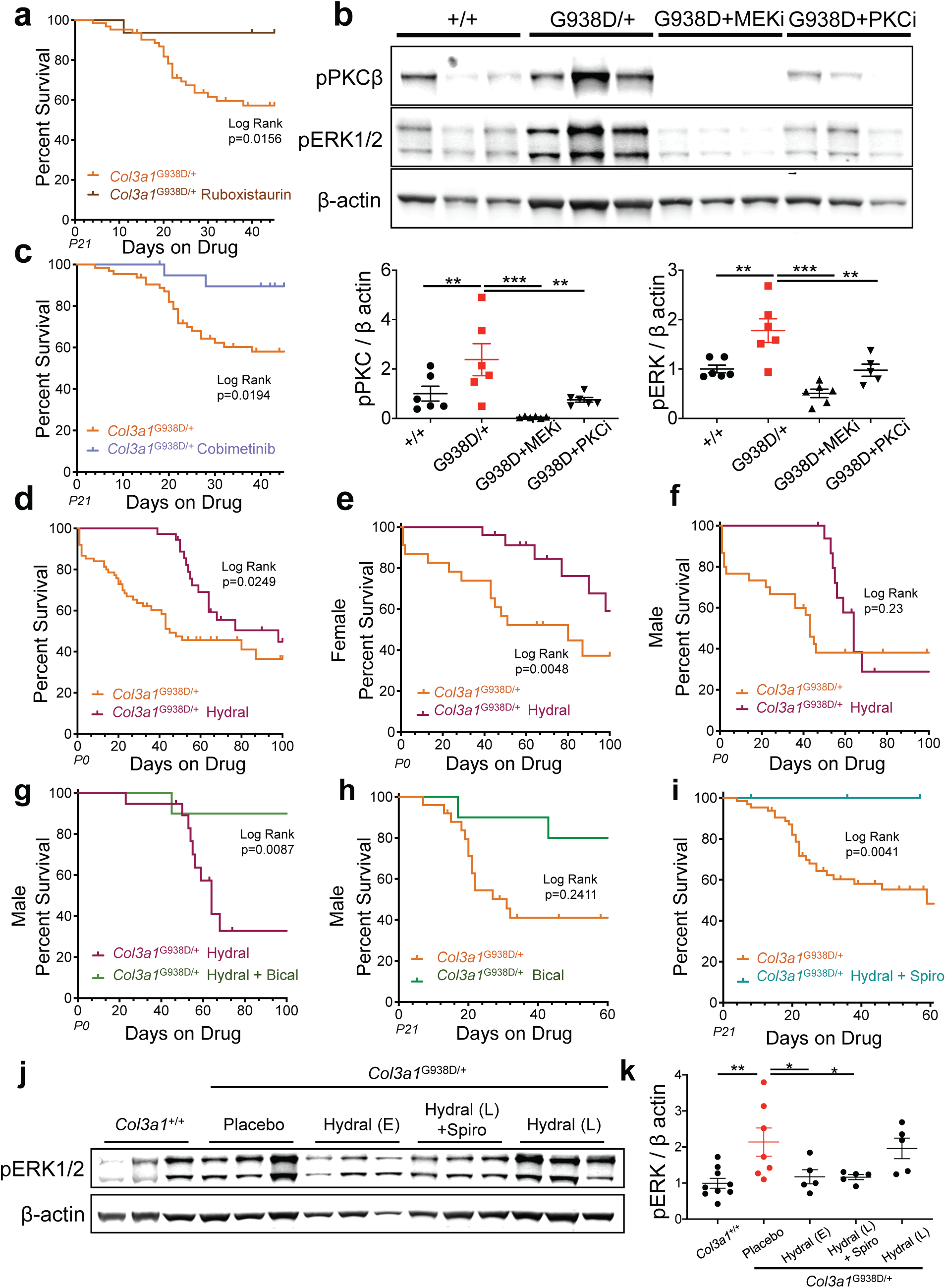
Inhibition of excessive PKC or ERK signaling prevents death due to aortic dissection. **a**, Kaplan-Meier survival curve comparing *Col3a1*^G938D/+^ (n=34) to *Col3a1*^G938D/+^ (n=16) mice receiving ruboxistaurin in the diet starting at weaning and continuing for 40 days. Significant differences were calculated using Log-Rank (Mantel-Cox) analysis. P21 = post-natal day 21 **b**, representative western blot analysis of pPKCβ and pERK comparing *Col3a1*^+/+^ (n=6) to *Col3a1*^G938D/+^ (n=6) proximal descending aortas and quantification of pPKCβ and pERK levels normalized to β-actin loading control for vEDS aortas. Error bars show mean ± s.e. Asterisks signify significant differences using one-way ANOVA with Dunnett’s multiple comparisons post-hoc test (**p<0.01, ***p<0.001, DF = 3, F = 13). **c**, Kaplan-Meier survival curve comparing *Col3a1*^G938D/+^ (n=34) to *Col3a1*^G938D/+^ (n=20) mice receiving cobimetinib in the drinking water starting at weaning and continuing for 40 days. Significant differences were calculated using Log-Rank (Mantel-Cox) analysis. P21 = post-natal day 21 **d**, Kaplan-Meier survival curve comparing *Col3a1*^G938D/+^ (n=43) to *Col3a1*^G938D/+^ (n=36) mice receiving hydralazine in the drinking water starting from birth and continuing for 3 months. Significant differences were calculated using Log-Rank (Mantel-Cox) analysis. Hydral = hydralazine; P0 = post-natal day 0 **e**, Kaplan-Meier survival curve comparing female *Col3a1*^G938D/+^ (n=23) to female *Col3a1*^G938D/+^ (n=26) mice receiving hydralazine in the drinking water starting from birth and continuing for 3 months. Significant differences were calculated using Log-Rank (Mantel-Cox) analysis. Hydral = hydralazine; P0 = post-natal day 0 **f**, Kaplan-Meier survival curve comparing male *Col3a1*^G938D/+^ (n=30) to male *Col3a1*^G938D/+^ (n=18) mice receiving hydralazine in the drinking water starting from birth and continuing for 3 months. Significant differences were calculated using Log-Rank (Mantel-Cox) analysis. Hydral = hydralazine; P0 = post-natal day 0 **g**, Kaplan-Meier survival curve comparing male *Col3a1*^G938D/+^ (n=18) mice receiving hydralazine in the drinking water starting from birth and continuing for 3 months to male *Col3a1*^G938D/+^ mice receiving hydralazine in the drinking water starting from birth and bicalutamide in the food starting from weaning and continuing for 2 months (n=10). Significant differences were calculated using Log-Rank (Mantel-Cox) analysis. Hydral = hydralazine; Bical = bicalutamide **g**, Kaplan-Meier survival curve comparing male *Col3a1*^G938D/+^ (n=30) mice male *Col3a1*^G938D/+^ mice receiving bicalutamide in the food starting from weaning and continuing for 2 months (n=10). Significant differences were calculated using Log-Rank (Mantel-Cox) analysis. Bical = bicalutamide; P21 = post-natal day 21 **i**, Kaplan-Meier survival curve comparing *Col3a1*^G938D/+^ (n=34) mice to *Col3a1*^G938D/+^ mice receiving hydralazine in the drinking water and spironolactone in the food starting from weaning and continuing for 2 months (n=16). Significant differences were calculated using Log-Rank (Mantel-Cox) analysis. Hydral = hydralazine; Spiro = spironolactone; P21 = post-natal day 21 **j**, representative western blot analysis of pERK comparing *Col3a1*^+/+^ (n=9) to *Col3a1*^G938D/+^ (n=7), *Col3a1*^G938D/+^ mice on hydralazine sampled at age P40 (Hydral (E), n=5), *Col3a1*^G938D/+^ mice on hydralazine and spironolactone sampled at age P70 (Hydral (L)+Spiro, n=5), and *Col3a1*^G938D/+^ on hydralazine sampled post-puberty at age P70 (Hydral (L), n=5) proximal descending aortas and quantification of pERK levels normalized to β-actin loading control for vEDS aortas. Error bars show mean ± s.e. Asterisks signify significant differences using one-way ANOVA with Dunnett’s multiple comparisons post-hoc test (**p<0.01, *p<0.05, DF = 4, F = 7.07).

In order to directly test the relevance of ERK activation we next treated mice with cobimetinib, an FDA-approved inhibitor of MEK, the kinase that activates ERK^21^. This treatment resulted in 90% survival after 45 days of treatment, compared to only 55% survival with no treatment (Fig. 3C). Immunoblot showed that increased survival correlated with the expected reduction in phosphorylation of ERK1/2, the downstream substrate for MEK but also, curiously, reduced PKC phosphorylation, suggesting the presence of a positive feedback loop (Fig. 3B). Neither cobimetinib nor ruboxistaurin had an effect on blood pressure (Supplementary Fig. 5).

In order to test if other FDA-approved medications that target the same pathway afford similar protection, we next treated our mice with hydralazine, a blood pressure medication that works, at least in part, by blocking IP_3_-mediated calcium release from the endoplasmic reticulum and hence PKC activation^22^. We observed improvement in survival, with 98% of treated mice surviving to 45 days – the median age of survival for an untreated vEDS mouse (Fig. 3D).

Protection from aortic dissection in vEDS mice treated with hydralazine was abruptly lost around the time of sexual maturity (∼50 days). While this was observed in both sexes, the effect was greatly exaggerated in male vs. female vEDS mice (25% vs. 60% survival at 100 days, respectively), suggesting a potential role for androgens (Fig. 3E, F). In keeping with this hypothesis, we found that the addition of the androgen receptor antagonist bicalutamide, at the time of weaning, to male mice that had been receiving hydralazine, lead to 90% survival at 100 days of age (Fig. 3G). Treatment with bicalutamide alone did not afford long-term protection from vascular dissection in vEDS mice (Fig. 3H). The elevated risk of vascular rupture in male mice at puberty is also seen in male vEDS patients and has been vaguely attributed to events that attend the adolescent growth spurt^4,23^. Incidence and severity of aneurysm/dissection is greater in males with MFS^24^ and males with abdominal aortic aneurysm^25^ suggesting that androgen signaling may play a broader role in aortic dissection, but the exact mechanism is still unclear^25–27^.

In anticipation of a reluctance to consider use of androgen receptor antagonists in young men with vEDS, we considered the potential use of spironolactone. Spironolactone is a diuretic that reduces adrenal androgen production, is a competitive antagonist of the androgen receptor and is used clinically to treat hyperandrogenism in the context of skin and hair disorders^28,29^. Spironolactone is commonly used in children of both genders for its diuretic effect and whereas males can experience mild gynecomastia and females can show menstrual irregularity, these effects are dose-and duration-dependent and are reversible with drug cessation. Initiation of treatment with hydralazine and spironolactone at 21 days of age in vEDS mice of both sexes achieved 100% survival at the end of the trial 60 days later, as compared to 45% survival in untreated animals (Fig. 3I). Survival correlated with the status of ERK1/2 phosphorylation in the aortic wall, which was increased in vEDS mice, reduced in vEDS mice prior to sexual maturity upon treatment with hydralazine, increased in vEDS mice after sexual maturity despite ongoing treatment with hydralazine, but again reduced in sexually mature vEDS mice treated with both hydralazine and spironolactone (Fig. 3J, K).

In women with vEDS, there is an estimated 12-25% lethality associated with pregnancy, historically attributed to hemodynamic stress^1,30,31^. We have previously shown that pregnancy-associated aortic dissection in MFS mice is largely driven by lactation-associated oxytocin release and oxytocin-induced PLC/IP_3_/PKC/ERK signaling^32^. In view of our data suggesting that over-activation of this pathway drives aortic dissection in vEDS mouse models, we hypothesized that pregnancy-associated vascular events in vEDS are driven by a mechanism similar to that observed in MFS mouse models.

We thus examined the effect of pregnancy and lactation on the survival of *Col3a1*^G209S/+^ mice, which have a milder phenotype. In this mouse model, we found that pregnancy and lactation is associated with 54% lethality due to arterial dissection in the first 30 days postpartum, compared to 96% survival during this same time period in never-pregnant *Col3a1*^G209S/+^ females (Fig. 4A). Removal of the litter right after birth and consequent prevention of lactation was able to prevent dissection and death in vEDS *Col3a1*^G209S/+^ mice, resulting in 100% survival in the postpartum period (Fig. 4B), suggesting that the increased risk of aortic dissection was dependent on lactation. Moreover, as previously shown in a mouse model of MFS^32^, treatment with a specific oxytocin receptor antagonist^33^ led to enhanced postpartum survival (∼90%) (Fig. 4C).

**Figure 4.**
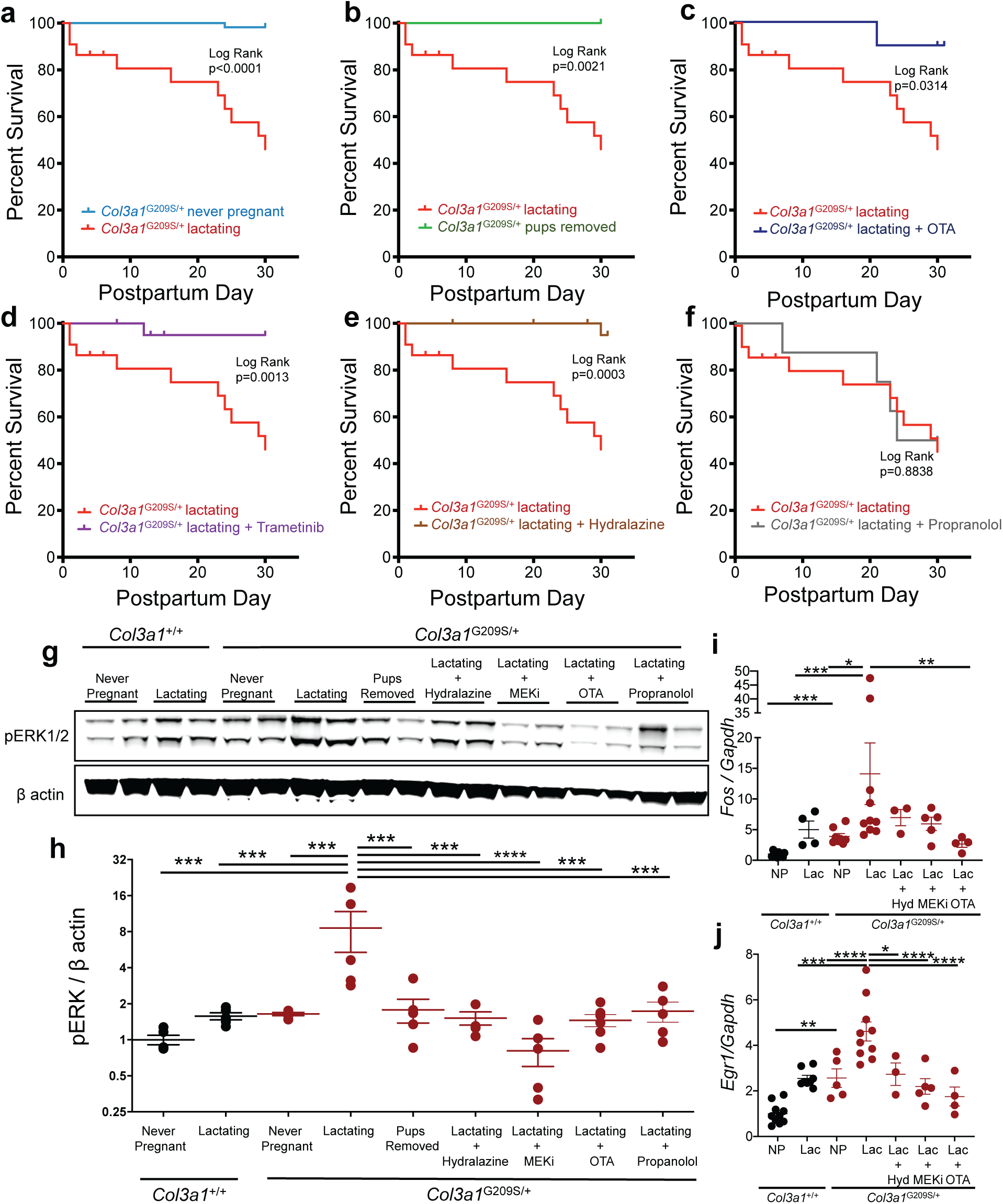
Oxytocin signaling during lactation increases the risk of aortic dissection. **a**, Kaplan-Meier survival curve for lactating *Col3a1*^G209S/+^ mice (n=22) compared to never pregnant female (n=55) *Col3a1*^G209S/+^ mice. Significant differences were calculated using Log-Rank (Mantel-Cox) analysis. **b**, Kaplan-Meier survival curve comparing *Col3a1*^G209S/+^ lactating (n=22) mice to *Col3a1*^G209S/+^ females with pups removed on the day of delivery thereby preventing lactation and eliminating the lactation-induced prolonged elevation of oxytocin (n=13). Significant differences were calculated using Log-Rank (Mantel-Cox) analysis. **c**, Kaplan-Meier survival curve comparing *Col3a1*^G209S/+^ lactating mice (n=22) to *Col3a1*^G209S/+^ females with oxytocin receptor antagonist (OTA) administered via a continuous subcutaneous infusion pump implanted at the end of the 3rd week of gestation and continued through the 4 weeks of lactation for a total of 5 weeks of treatment (n=10). Significant differences were calculated using Log-Rank (Mantel-Cox) analysis. **d**, Kaplan-Meier curve demonstrating the survival of *Col3a1*^G209S/+^ lactating mice treated with trametinib (n=21), a MEK inhibitor, initiated at the start of the 3rd week of pregnancy and continued through 4 weeks of lactation, in comparison to untreated lactating *Col3a1*^G209S/+^ mice (n=22). Significant differences were calculated using Log-Rank (Mantel-Cox) analysis. **e**, Kaplan-Meier curve demonstrating the survival of *Col3a1*^G209S/+^ lactating mice treated with hydralazine (n=23) initiated at the start of the 3rd week of pregnancy and continued through 4 weeks of lactation, in comparison to lactating untreated *Col3a1*^G209S/+^ mice (n=22). Significant differences were calculated using Log-Rank (Mantel-Cox) analysis. **f**, Kaplan-Meier curve demonstrating the survival of *Col3a1*^G209S/+^ lactating mice treated with propranolol (n=8), initiated at the start of the 3rd week of pregnancy and continued through 4 weeks of lactation, in comparison to lactating untreated *Col3a1*^G209S/+^ mice (n=22). Significant differences were calculated using Log-Rank (Mantel-Cox) analysis. **g**, Representative western blot analysis of pERK1/2 in the proximal descending aorta comparing *Col3a1*^+/+^ never pregnant, *Col3a1*^+/+^ lactating, *Col3a1*^G209S/+^ never pregnant, *Col3a1*^G209S/+^ lactating mice, *Col3a1*^G209S/+^ mice with pups removed thereby preventing the lactation, and *Col3a1*^G209S/+^ mice treated with trametinib (MEKi) or oxytocin receptor antagonist (OTA) or hydralazine or propranolol. **h**, Quantification of western Blot analysis of pERK1/2 in the proximal descending aorta in *Col3a1*^+/+^ never pregnant (n=5), *Col3a1*^+/+^ lactating (n=5), *Col3a1*^G209S/+^ never pregnant (n=5), *Col3a1*^G209S/+^ lactating mice (n=5), *Col3a1*^G209S/+^ mice with pups removed thereby preventing the lactation (n=5), and *Col3a1*^G209S/+^ mice treated with trametinib (n=5, MEKi) or oxytocin receptor antagonist (n=5, OTA) or hydralazine (n=3) or propranolol (n=5). Error bars show mean ± s.e. Asterisks signify significant differences of log-transformed data using one-way ANOVA with Dunnett’s multiple comparisons post-hoc test (*p<0.05, **p<0.01, ***p<0.001, ****p<0.0001, DF = 8, F = 8.06) **i**, qPCR analysis of *Fos* normalized to *Gapdh* in the proximal descending aorta comparing wild-type never pregnant (n=6), wild-type lactating (n=4), *Col3a1*^G209S/+^ never pregnant (n=8), *Col3a1*^G209S/+^ lactating mice (n=10), and *Col3a1*^G209S/+^ mice treated with trametinib (n=5, MEKi) or oxytocin receptor antagonist (n=4, OTA) or hydralazine (n=3, Hyd). Error bars show mean ± s.e. Asterisks signify significant differences of log-transformed data using one-way ANOVA with Dunnett’s multiple comparisons post-hoc test (*p<0.05, **p<0.01, ***p<0.001 DF = 6, F = 13.84) **j**, qPCR analysis of *Egr1* normalized to *Gapdh* in the proximal descending aorta comparing *Col3a1*^+/+^ never pregnant (n=10), *Col3a1*^+/+^ lactating (n=7), *Col3a1*^G209S/+^ never pregnant (n=5), *Col3a1*^G209S/+^ lactating mice (n=10), and *Col3a1*^G209S/+^ mice treated with trametinib (n=5, MEKi) or oxytocin receptor antagonist (n=4, OTA) or hydralazine (n=3, Hyd). Error bars show mean ± s.e. Asterisks signify significant differences using one-way ANOVA with Dunnett’s multiple comparisons post-hoc test (*p<0.05, **p<0.01, ***p<0.001, ****p<0.0001 DF = 6, F = 14.84)

We next tested the effect of inhibitors of MEK/ERK and PKC activation in postpartum lactating female mice. Initiation of either MEK/ERK inhibition using trametinib or IP_3_/PKC inhibition using hydralazine in *Col3a1*^G209S/+^ mice at the time of delivery resulted in 95% survival at day 30 postpartum, compared to 46% survival in untreated lactating vEDS mice (Fig. 4D, E). The beneficial effect of these treatments correlated with a decrease in ERK activation, as measured immunoblotting of aortic lysates for ERK1/2 phosphorylation and analysis of gene expression of ERK target genes (Fig. 4G-J). Similar to what we observed in non-pregnant vEDS mice (Supplementary Fig. 3), treatment with propranolol had no impact on survival in the postpartum period (Fig. 4F). This suggests that despite the reduction in blood pressure and heart rate achieved with propranolol (Supplementary Fig. 3), a current standard of care for pregnant women with vEDS, there was no demonstrable beneficial effect on survival in this mouse model. Taken together, this data further supports the notion that excessive activation of the PLC/IP_3_/PKC/ERK signaling pathway drives aortic dissection in both pregnant and non-pregnant vEDS mice.

Our results provide the first evidence for a targetable signaling abnormality that contributes to the pathogenesis of vEDS and illustrate the promise of therapeutic strategies aimed at inhibition of the PLC/IP_3_/PKC/MEK/ERK axis of activation. It is notable that other vascular connective tissue disorders converge on the same pathway despite mechanistic differences in more proximal events (e.g. TGFβ and AT1R activation in MFS mouse models). Additional work will be needed to elucidate the nature of outside-in cellular signaling that initiates with type III collagen deficiency, cross-talks with androgen signaling, and culminates in PKC/ERK activation and arterial dissection in vEDS.

## Methods

### Mice

All mice were cared for under strict adherence to the Animal Care and Use Committee of the Johns Hopkins University School of Medicine.

#### Generation of Col3a1 knock-in mice

To generate *Col3a1* knock-in mice, we targeted the *Col3a1* locus using CRISPR–Cas9 (clustered, regularly interspaced palindromic repeat–CRISPR-associated protein 9). Two single-guide RNA (sgRNA) sequences for each intended mutation (see Supplementary Table 1) were designed to target exon 39 or exon 7 of *Col3a1* (NC_000067.6) using a gRNA CRISPR design tool (crispr.mit.edu). The gRNAs were predicted to have negligible off-target effects. The sgRNA was transcribed in vitro. The homology directed repair (HDR) template was purchased as a 4-nmol Ultramer (IDT, see Supplementary Table 1). For the *Col3a1*^G938D/+^ mouse, the sgRNA, Cas9 (TriLink BioTechnologies), and HDR were co-injected into C57BL/6J zygotes (Johns Hopkins University Transgenic Core). For the *Col3a1*^G209S/+^ mouse, the guide RNA sequences were ligated into the pX330 vector (Addgene#42230) according to Cong et al.^34^. Briefly, plasmid was linearized with BbsI (R3539S, New England Biolabs) and gRNAs were ligated into the restriction site and verified through Sanger sequencing. In vitro transcription of the cloned gRNAs was performed using MEGAscript T7 Transcription Kit (Life Technologies). The amplicon was purified with a PCR purification kit (Qiagen) and was used as a template for the IVT kit, according to the suggested protocol. All mice were maintained on a C57BL/6J background (#000664, The Jackson Laboratory).

#### Mutation validation, Sanger sequencing, and Genotyping

Bidirectional Sanger DNA sequencing assays were performed using primers designed 100 to 200 bp from the intended mutation to confirm correct insertion. PCR was performed using a DNA Engine Dyad thermal cycler (Bio-Rad). iProof High Fidelity PCR Master Mix (Bio-Rad) was used in accordance with the manufacturer’s instructions for each primer set. Cycle sequencing was performed using the BigDye Terminator v3.1 kit and an ABI 3730xl DNA Analyzer in accordance with the manufacturer’s instructions (Life Technologies). Samples were purified using the QIAquick PCR Purification kit (Qiagen). After confirmation of the intended mutation, restriction enzymes were used to detect the presence or absence of the mutation. The G209S mutation leads to the loss of an AvaII cut site and the G938D mutation leads to the gain of a BamHI cut site (AvaII R0153L; BamHI-HF R3136S, New England Biolabs).

All mice found dead were assessed for cause of death by necropsy, noting in particular hemothorax and hemoperitoneum.

### Echocardiography

Mice were imaged as previously described^12^. Briefly, echocardiograms were performed on awake, unsedated mice using the Visualsonics Vevo 660 V1.3.6 imaging system and a 30MHz transducer. Three separate measurements of the maximal internal dimension at the aortic root and proximal ascending aorta were made from distinct captured images and averaged. All imaging and measurements were performed by a researcher who was blinded to genotype. Measurements were normalized to weight, as the *Col3a1*^G938D/+^ mice are significantly smaller than their *Col3a1*^+/+^littermates.

### Histology

Mice were euthanized by isoflurane inhalation and the left common iliac artery was transected to allow for drainage. PBS (pH 7.4) and PBS containing 4% paraformaldehyde (PFA) was flushed through the left ventricle. The heart and thoracic aorta were removed en block and fixed in 4% PFA overnight at 4°C. Aortas were submitted for paraffin fixation and longitudinal sections 5 micrometers thick were mounted on glass slides and stained with hematoxylin & eosin (HE), Verhoeff-van Giesen (VVG), Masson’s Trichrome, or Picrosirius red (PSR). Slides were imaged at 20x and 40x magnification using a Nikon Eclipse E400 microscope. Collagen content was determined by polarized PSR intensity^35^ and elastin breaks were counted by a researcher blinded to genotype and treatment arm.

### Electron Microscopy

Electron microscopy was performed as previously described^36^ focusing on the proximal descending thoracic aortic wall.

### RNAseq

RNA was isolated from the proximal descending thoracic aorta of three mice for each condition, flushed in PBS, and directly stored into TRIzol (Invitrogen). RNA was extracted according to manufacturer’s instructions and purified using the PureLink RNA Mini Kit (Invitrogen). Library prep was performed using TruSeq Stranded Total RNA with Ribo-Zero (Illumina). Sequencing was run on an Illumina HiSeq2500 using standard protocols.

### Bioinformatics

Illumina’s CASAVA 1.8.4 was used to convert BCL files to FASTQ files. Default parameters were used. rsem-1.3.0 was used for running the alignments as well as generating gene and transcript expression levels. The data was aligned to “mm10” reference genome. EBseq was used for Differential Expression analysis and default parameters were used^37^. The networks and upstream regulator analyses were generated through the use of IPA (QIAGEN Inc., https://www.qiagenbioinformatics.com/products/ingenuity-pathway-analysis).

### Western Blot

Mouse descending thoracic aortas (distal to the left subclavian branch) were harvested, snap frozen in liquid nitrogen, and stored at −80°C until processed. Protein was extracted using an automatic bead homogenizer in conjunction with a Protein Extraction Kit (Full Moon Biosystems). All protein lysis buffers contained both PhosSTOP and cOmplete™, Mini, EDTA-free Protease Inhibitor Cocktail (Roche). Western blotting was performed using LI-COR buffer and species appropriate secondary antibodies conjugated to IR-dye700 or IRdye-800 (LI-COR Biosciences), according to the manufacturer’s guidelines and analyzed using LI-COR Odyssey. The following primary antibodies were used: anti-β-Actin (8H10D10) (Cell Signaling Technology, 3700), anti-phospho ERK1/2 (Cell Signaling Technology, 4370), anti-PKC (phospho S660) (Abcam, 75837).

### Delivery of Medication

For drug trials in the G938D mutation, mice were initiated on medication at weaning and continued until 2 months of age. Cobimetinib (GDC-0973/RO551404, Active Biochem) was dissolved in drinking water and filtered to reach a final concentration of 0.02g/L giving an estimated dose of 2mg/kg/day. Ruboxistaurin (LY333531 HCl, Selleck Chemicals) was mixed with powdered food (LabDiet) to give a concentration of 0.1mg/g giving an estimated dose of 8 mg/kg/day. Hydralazine (Exelan) is known to be safe in pregnancy and is found in breastmilk, so was initiated at birth and dissolved in drinking water and filtered to reach a final concentration of 0.32g/L, giving an estimated daily dose of 32mg/kg/day. Bicalutamide (Major Pharmaceuticals) was mixed with powdered food to give a concentration of 0.625mg/g giving an estimated dose of 50mg/kg/day. Losartan (Merck) was dissolved in drinking water and filtered to reach a final concentration of 0.6g/L giving an estimated dose of 60mg/kg/day. Celiprolol (Pfizer) was dissolved in drinking water and filtered to reach a final concentration of 2g/L giving an estimated dose of 200mg/kg/day. Propranolol (Qualitest) was dissolved in the drinking water and filtered to reach a final concentration of 0.8g/L giving an estimated dose of 80mg/kg/day. Atenolol (Teva) was dissolved in the drinking water and filtered to reach a final concentration of 1.2g/L giving an estimated dose of 120mg/kg/day. Amlodipine (Zygenerics) was dissolved in drinking water and filtered to reach a final concentration of 0.12g/L giving an estimated dose of 12mg/kg/day. Spironolactone (Sky) was mixed with powdered food (LabDiet) to give a concentration of 1.25mg/g giving an estimated dose of 100mg/kg/day.

For the drug trials in pregnancy, mice were initiated on medication at the 3^rd^ week of gestation and continued for 1 month postpartum. The selective oxytocin antagonist (des Gly-NH2,d(CH2)5[D-Tyr2,Thr4]OVT) was synthesized in the laboratory of Dr. Maurice Manning (University of Toledo) and was delivered in phosphate buffered saline (PBS) to reach a final dose of 1μg/kg/hr when delivered via continuous infusion in a mini Alzet pump implanted subcutaneously between the scapulae. Propranolol (Qualitest) was dissolved in the drinking water and filtered to reach a final concentration of 0.8g/L giving an estimated dose of 80mg/kg/day. Hydralazine (Exelan) was dissolved in drinking water and filtered to reach a final concentration of 0.16g/L, giving an estimated daily dose of 16mg/kg/day. Trametinib (GSK) was dissolved in PBS with 10% DMSO and mice were treated once a day by oral gavage, giving an estimated daily dose of 1mg/kg/day. Placebo-treated animals received regular drinking water.

### Blood Pressure Analysis

Blood pressures were measured by tail cuff plethysmography one week prior to completion of a study. Mice were habituated to the system prior to collection in which 10-15 measurements were obtained and averaged.

### Statistical Analysis

All data points are presented for quantitative data, with an overlay of the mean with SEM or as interleaved box & whiskers showing the minimum to maximum. All statistical analysis was performed using GraphPad Prism 8.

For data that did not pass Shapiro-Wilk normality tests, Kruskal-Wallis (nonparametric) tests were performed to evaluate significance between groups using Dunn’s multiple comparison test with a p-value of <0.05 considered statistically significant. For data that did pass normality, two-way ANOVA was used with multiple comparisons.

For single comparisons, if the Shapiro-Wilk normality test was passed, then two-tailed unpaired t-tests were performed. If Shapiro-Wilk normality test did not pass, then Mann-Whitney nonparametric tests were performed.

Kaplan-Meier survival curves were compared using a log-rank (Mantel-Cox) test. While each treatment trial included contemporaneous control animals, the performance of untreated mice remained constant for the full duration of this study, allowing pooling of controls to improve statistical power.

## Acknowledgements

We thank the staff of the Next Generation Sequencing Center, Research Animal Resources, and the Transgenic Core Laboratory at Johns Hopkins University School of Medicine for their assistance with various aspects of this project. We thank Djahida Bedja (Johns Hopkins University) for her assistance with echocardiograms and Maurice Manning (University of Toledo) for providing the oxytocin receptor antagonist and guidance regarding its use in animal studies.

Supported by grants from the National Institutes of Health (AR41135 to H.C.D.; GM007309 to C.J.B.), the Howard Hughes Medical Institute (to H.C.D.), the Marfan Foundation (to H.C.D.), the Ehlers Danlos Syndrome Network C.A.R.E.S. Foundation (to H.C.D.), EDS Today (to H.C.D.), the Daskal Family Foundation (to H.C.D.), the Alison Aldredge Family Foundation (to H.C.D.), the DEFY Foundation (to H.C.D.), the Natural Sciences and Engineering Research Council of Canada (to E.C.D.), the Fulbright-Conicyt Scholarship (to J.F.C.G.), and Fondecyt Grant (11170353 to J.F.C.G.).

## Author contributions

C.J.B. and H.C.D. were responsible for all aspects of study design, data interpretation and preparation of the manuscript. C.J.B., Z.B., and G.R. were responsible for mouse husbandry and medication dosing. C.J.B., J.F.C.G., G.R., and M.H. created and phenotyped mouse models. E.C.D. performed electron microscopy studies. C.J.B. performed and interpreted echocardiograms. C.J.B., E.G.M., and H.C.D. wrote the manuscript with contributions from all authors.

## Data availability

RNA sequencing data is available from the Sequence Read Archive (SRA) under accession PRJNA532935. All other data associated with this study are present in the paper or supplementary materials.

**Supplementary Table 1.** List of primers and oligonucleotide sequences used in the *Col3a1*^G209S/+^ and *Col3a1*^G938D/+^ mouse studies.

**Supplementary Figure 1.** vEDS mice do not have ascending aortic aneurysm.

**a**, Echocardiographic images of the root and ascending aorta in wild-type and vEDS mice. Yellow arrowheads point to the root and green arrowheads point to the ascending aorta.

**b**, Weights of male and female vEDS mice at 45 days of age. Error bars show mean ± s.e. Asterisks signify significant differences using two-way ANOVA (***p<0.001; DF = 2, F = 0.3943 interaction; DF = 1, F = 17.18 sex; DF = 2, F = 10.42 genotype)

**c**, Measurement of aortic root diameter normalized to mouse weight. Error bars show mean ± s.e. There are no significant differences using Kruskal-Wallis with Dunn’s multiple comparisons post-hoc test.

**d**, Measurement of ascending aortic diameter normalized to mouse weight. Error bars show mean ± s.e. There are no significant differences using Kruskal-Wallis with Dunn’s multiple comparisons post-hoc test.

**Supplementary Figure 2.** vEDS aortas have abnormal extracellular matrix architecture.

**a-c**, electron microscopy images of the proximal descending aorta in wild-type and vEDS mice. SMC = vascular smooth muscle cell, e = elastin, col = collagen fibril

**d-f**, increased magnification of elastin-collagen-vascular smooth muscle cell interface in *Col3a1*^+/+^, *Col3a1*^G209S/+^, and *Col3a1*^G938D/+^ aortas.

**g-i** electron microscopy images of collagen cross fiber diameter in *Col3a1*^+/+^, *Col3a1*^G209S/+^ and *Col3a1*^G938D/+^ aortas.

**j**, density plot demonstrating the distribution of collagen cross fiber diameters for *Col3a1*^+/+^, *Col3a1*^G209S/+^, and *Col3a1*^G938D/+^ aortas.

**k**,**l** electron microscopy images of adventitial fibroblasts in *Col3a1*^+/+^ and *Col3a1*^G938D/+^ aortas. ER = endoplasmic reticulum

**Supplementary Figure 3.** Losartan, propranolol, atenolol, amlodipine, and celiprolol do not rescue death from aortic dissection.

**a**, Kaplan-Meier survival curve comparing *Col3a1*^G938D/+^ (n=34) to *Col3a1*^G938D/+^ (n=24) mice receiving losartan in the drinking water. Significant differences were calculated using Log-Rank (Mantel-Cox) analysis. P21 = post-natal day 21

**b**, Kaplan-Meier survival curve comparing *Col3a1*^G938D/+^ (n=34) to *Col3a1*^G938D/+^ (n=9) mice receiving propranolol in the drinking water. Significant differences were calculated using Log-Rank (Mantel-Cox) analysis. P21 = post-natal day 21

**c**, Kaplan-Meier survival curve comparing *Col3a1*^G938D/+^ (n=34) to *Col3a1*^G938D/+^ (n=13) mice receiving atenolol in the drinking water. Significant differences were calculated using Log-Rank (Mantel-Cox) analysis. P21 = post-natal day 21

**d**, Kaplan-Meier survival curve comparing *Col3a1*^G938D/+^ (n=34) to *Col3a1*^G938D/+^ (n=11) mice receiving amlodipine in the drinking water. Significant differences were calculated using Log-Rank (Mantel-Cox) analysis. P21 = post-natal day 21

**e**, Kaplan-Meier survival curve comparing *Col3a1*^G938D/+^ (n=34) to *Col3a1*^G938D/+^ (n=18) mice receiving celiprolol in the drinking water. Significant differences were calculated using Log-Rank (Mantel-Cox) analysis. P21 = post-natal day 21

**f**, Kaplan-Meier survival curve comparing *Col3a1*^G209S/+^ (n=12) to *Col3a1*^G209S/+^ (n=11) mice receiving celiprolol in the drinking water. Significant differences were calculated using Log-Rank (Mantel-Cox) analysis. P60 = post-natal day 60

**g**, blood pressure measurements for mice on losartan. Error bars show mean ± s.e. Asterisks signify significant differences between treatment groups using two-way ANOVA for blood pressure (DF = 1, F=20.39) and Student’s t-test for pulse rate (t=0.5155)

**h**, blood pressure measurements for mice on propranolol. Error bars show mean ± s.e. Asterisks signify significant differences between treatment groups using two-way ANOVA for blood pressure (DF = 1, F =15.26) and Mann-Whitney test for pulse rate (**p<0.01).

**i**, blood pressure measurements for mice on atenolol. Error bars show mean ± s.e. Asterisks signify significant differences between treatment groups using two-way ANOVA for blood pressure (DF=1, F=5.90) and Student’s t-test for pulse rate (****p<0.0001, t=20.12)

**j**, blood pressure measurements for mice on amlodipine. Error bars show mean ± s.e. Asterisks signify significant differences between treatment groups using two-way ANOVA for blood pressure (DF=1, F=8.134) and Student’s t-test for pulse rate (p=0.1511, t=1.569)

**k**, blood pressure measurements for mice on celiprolol. Error bars show mean ± s.e. Asterisks signify significant differences between treatment groups using two-way ANOVA for blood pressure (DF=1, F=272.8) and Student’s t-test for pulse rate (**p<0.01, t=3.959)

**Supplementary Figure 4.** Pathway analysis (Ingenuity) based on the differentially expressed genes identified by bulk RNAseq.

**Supplementary Figure 5.** Cobimetinib and Ruboxistaurin do not lower blood pressure.

**a**, blood pressure measurements for mice on cobimetinib. Error bars show mean ± s.e. Differences between treatment groups were calculated two-way ANOVA for blood pressure and Student’s t-test for pulse rate (DF=1, F=0.7523, T=3.325)

**b**, blood pressure measurements for mice on ruboxistaurin. Error bars show mean ± s.e. Differences between treatment groups were calculated using two-way ANOVA for blood pressure and Student’s t-test for pulse rate (DF = 1, F=0.0179, T=2.488)

**c**, blood pressure measurements for mice on hydralazine. Error bars show mean ± s.e. Differences between treatment groups were calculated using two-way ANOVA for blood pressure and Student’s t-test for pulse rate (DF = 1, F=6.810, T=3.421)

