## Supplemental Table 1 for "Targetable cellular signaling events drive arterial rupture in knock-in mouse models of vascular Ehlers-Danlos Syndrome"

Supplementary Table 1. List of primers and oligonucleotide sequences used in the G938D/+ and G209S/+ mice studies

| *Col3a1* Primer | Forward | Reverse |
|---|---|---|
| G209S | TTTACATTCCAGGGTCCACCAG | GTCAGGAATCTCTAAAGTTCACCTC |
| G938D | TCCTGGTCCTCCTGGCAATA | TCAGATGCAAGGTGACTATGCT |
| gRNA Target Site | gRNA1 | gRNA2 |
| Exon 7 | CCTGGTGAACCTGGTCAAGC | CCAGGGGGACCTTGGTATC |
| Exon 39 | AGTGGTGCTCCTGGCAAGGA | ACAGCAATAGCTCTCACCGG |
| HDR Template | | |
| Exon 7 Target | TATATTTACACTATTCTTCTATTTTAGGGTTCTCCTGGATACCAAGGTCCCCCTGGTGAACCTGGTCAAGCTTCTCCAGCA GTAAGTAACACTTGAGAATTGCTAAACACATTG | |
| Exon 39 Target 1 | GGCAAGGAAAGCAAAAAGACCCTTTTCTGCCATGTTAAAATTTAAATTATTTGCAGGGTAATCCAGGGCCCCCAGGACCCAGTGGTGCTCCTGGCAAGGACGGCCCTCCAGGTCCTGCAGGCAACAGTGGTTCTCCTGGCAACCCTGGAATAGCTGGACCAAAAGGTGATGCTGGACAGCCTGGAGAGAAGGGGCCACCTGGTGCTCAG(G/A)TCCTCCGGTGAGAGCTATTGCTGTTTTGTTGGTAGACTGCATATTCTGATAACAAACATAATAAGGGAGCTAGGATTCTGTAGCCATAGGCTGATTCTTT | |
| Exon 39 Target 2 | TCCTGCAGGCAACAGTGGTTCTCCTGGCAACCCTGGAATAGCTGGACCAAAAGGTGATGCTGGACAGCCTGGAGAGAAGGGGCCACCTGGTGCTCAG(G/A)TCCTCCGGTGAGAGCTATTGCTGTTTTGTTGGTAGACTGCATATTCTGATAACAAACATAATAAGGGAGCTAGGATTCTGTAGCCATAGGCTGATTCTTTC | |
