## Supplementary figures and images for "Targetable cellular signaling events drive arterial rupture in knock-in mouse models of vascular Ehlers-Danlos Syndrome"

### Supplemental Figure 1

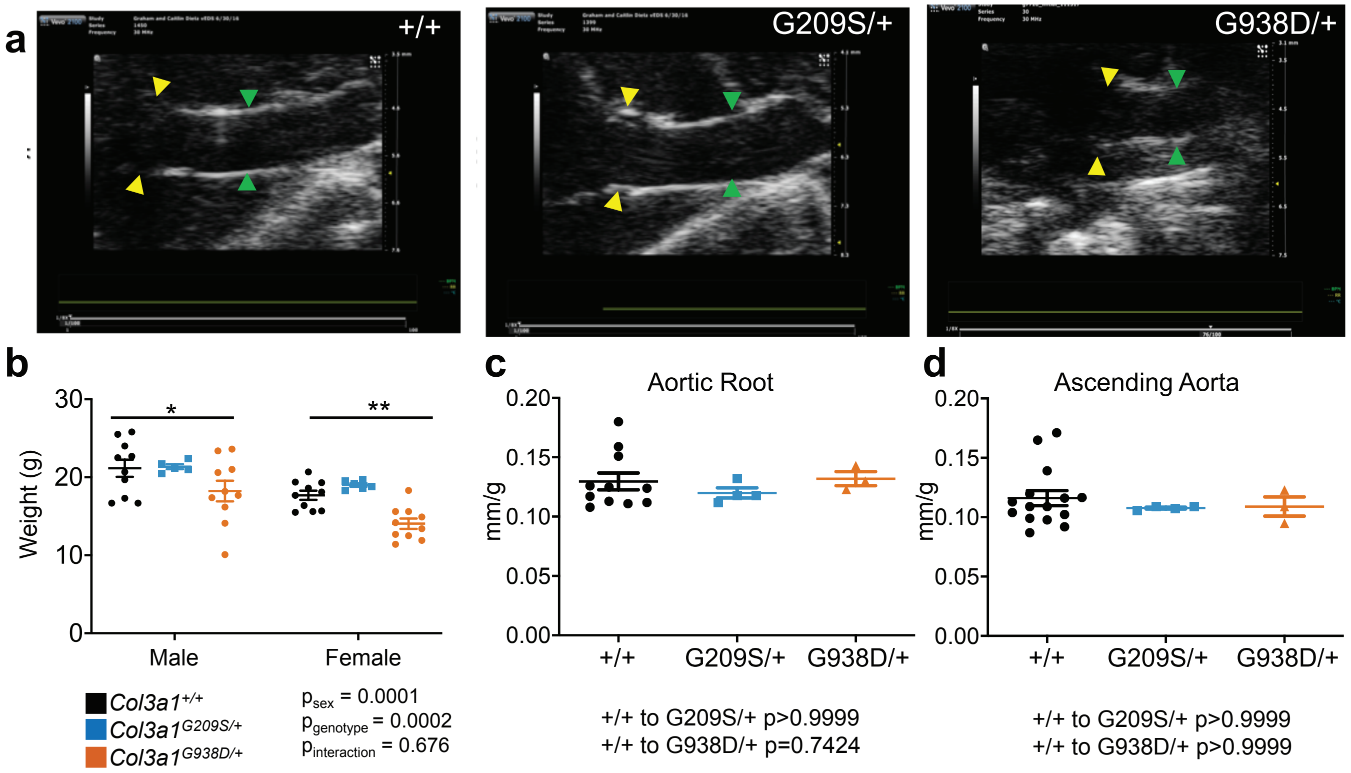


Supplementary Figure 1.

### Supplemental Figure 2

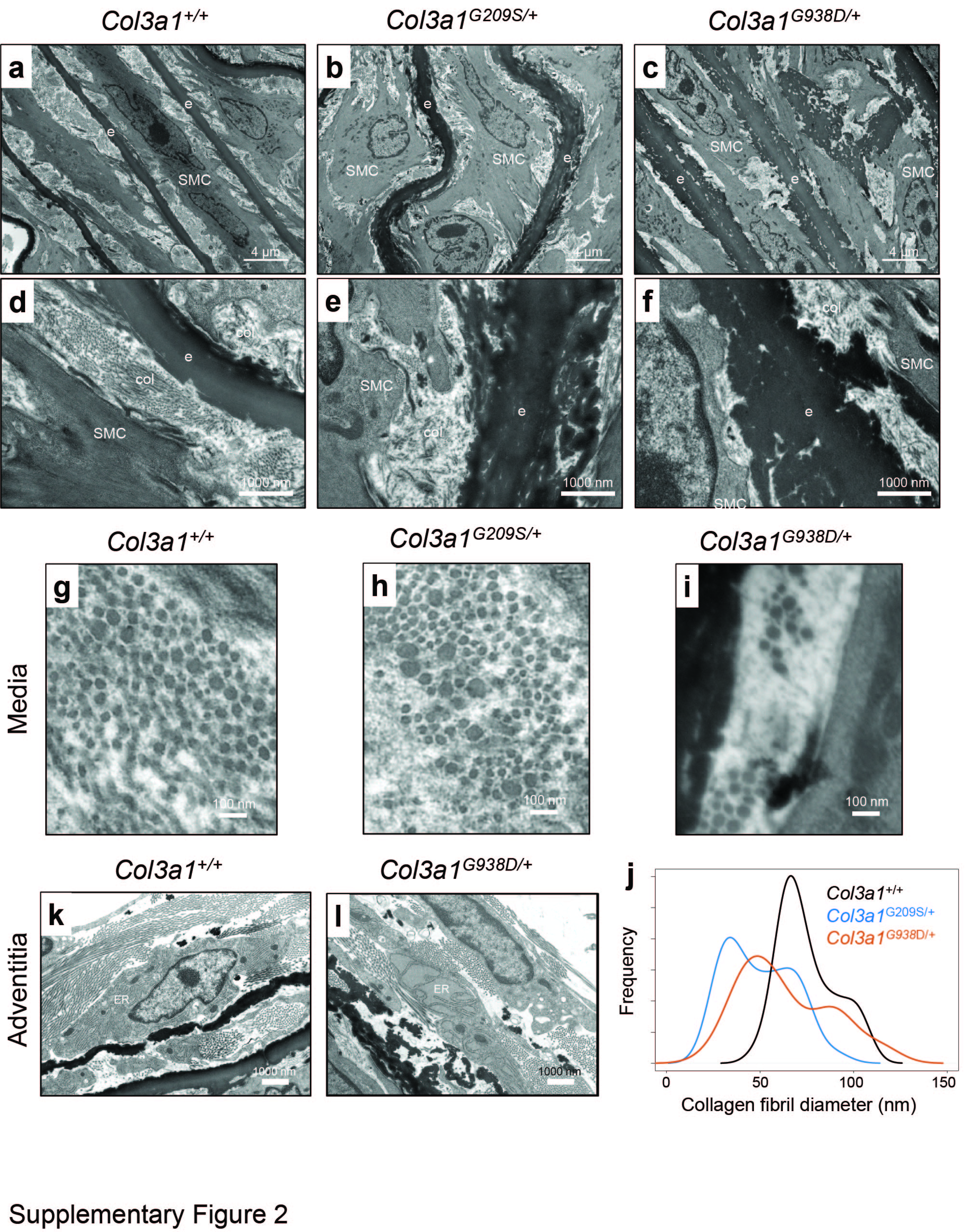

### Supplemental Figure 3

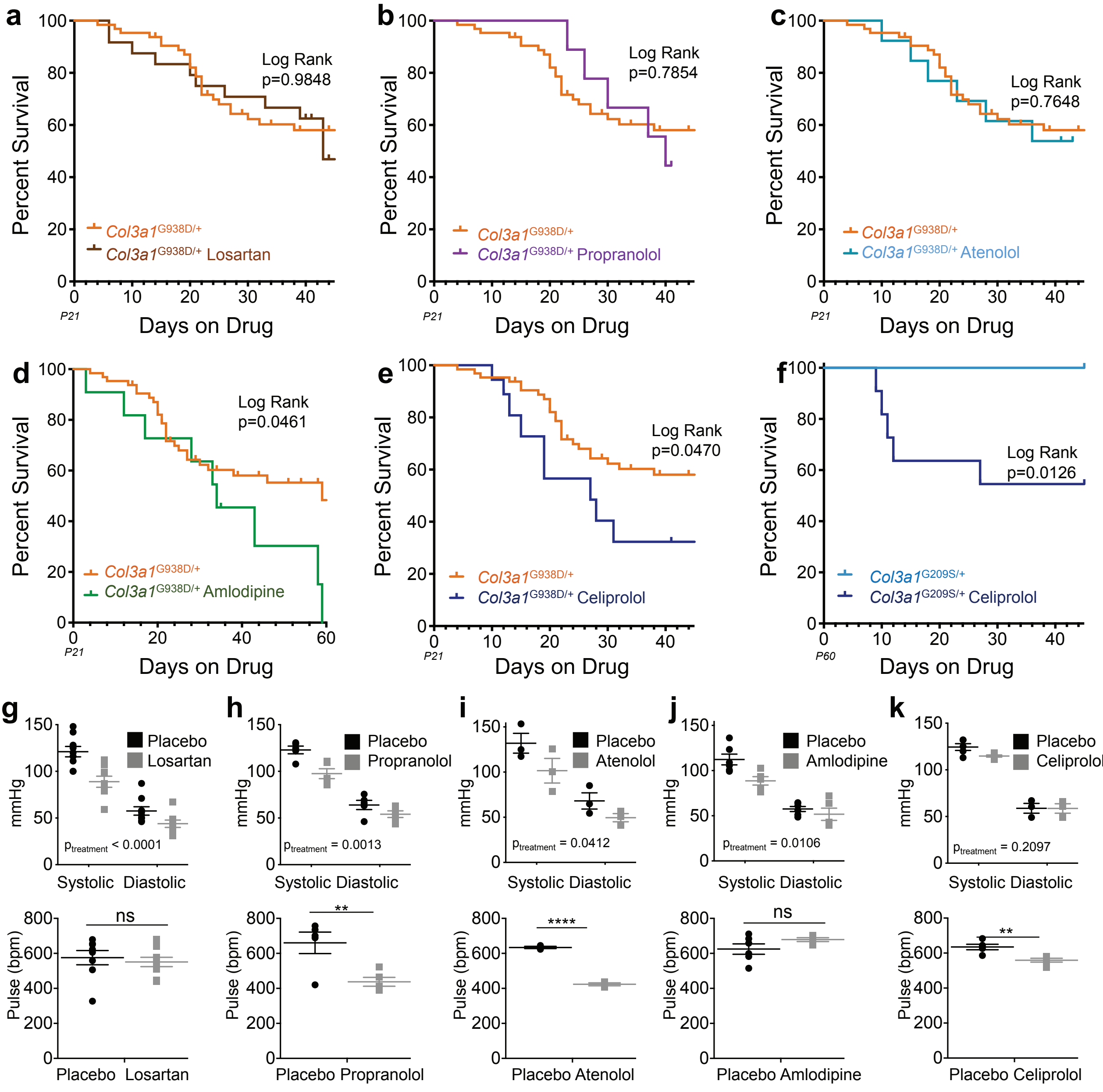


Supplementary Figure 3.

### Supplemental Figure 4

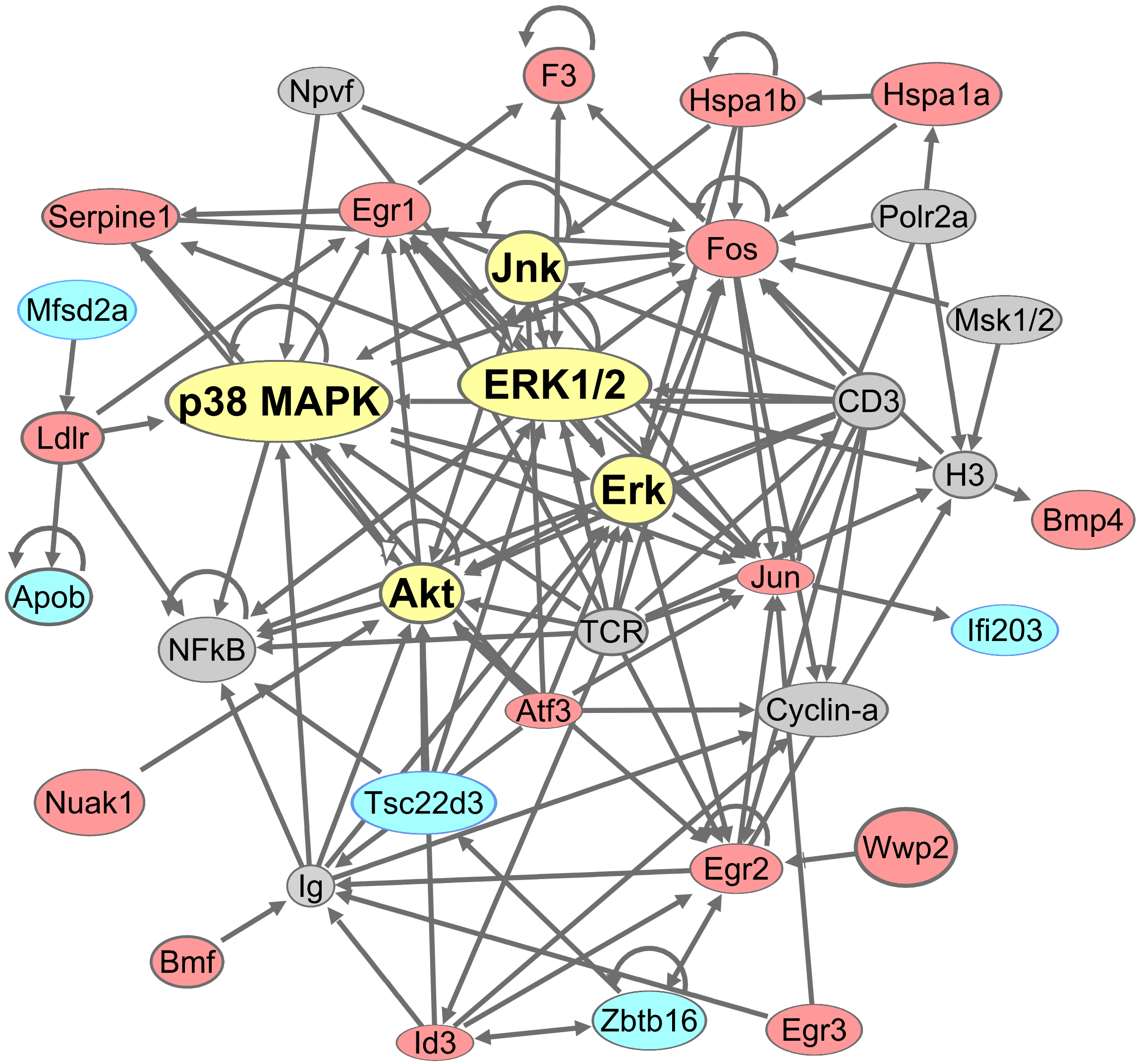


Supplementary Figure 4

### Supplemental Figure 5

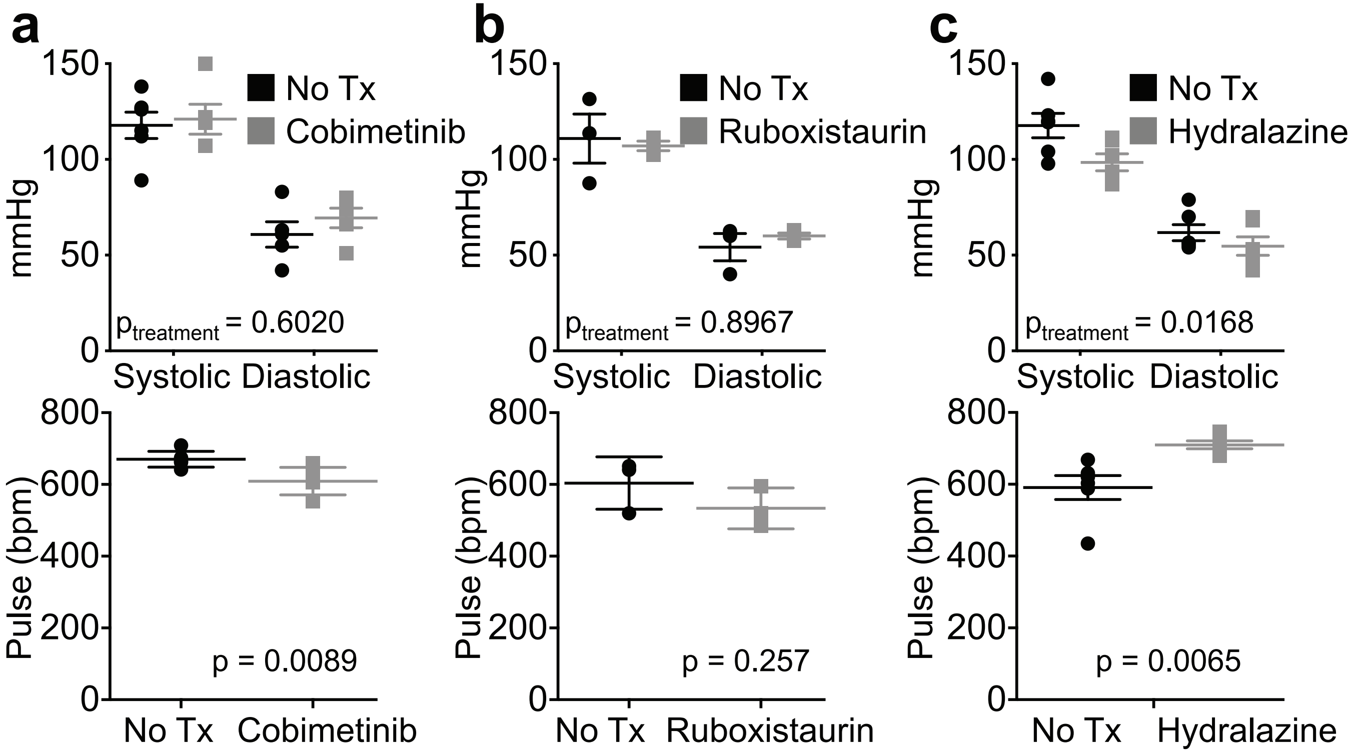


Supplementary Figure 5
